## Supplementary Information for "The spread of the first introns in proto-eukaryotic paralogs"

**Supplementary Text**

*The emergence of two different intron types*

Two types of spliceosomal introns emerged during eukaryogenesis: U2 and U12. Three different models for the appearance of two types have been proposed (Burge et al. 1998). The first is the codivergence model, which postulates that the snRNA genes and introns diverged into two different sets after duplication of the snRNA genes. According to the fission/fusion model the two intron types evolved in separate proto-eukaryotic lineages that later fused. Whereas in the first two models the two types originated from primordial spliceosomal introns, in the parasitic invasion model the two types represent two temporally separate invasions of self-splicing group II introns.

To investigate the origin of the two different types of introns during eukaryogenesis, we predicted the type of all introns. 1.5% (KOGs) and 1.9% (Pfams) of LECA introns were predicted to be of U12-type, which was higher than in any present-day eukaryote in our dataset. 14.8% (KOGs) and 31.1% (Pfams) of these U12-type LECA introns were shared with an intron in a paralog, which is not significantly lower than for U2-type introns (Fisher's exact tests, $P = 0.072$ (KOGs) and $P = 0.53$ (Pfams)). Most of these shared U12-type LECA introns in KOGs were paired to U2-type introns in paralogs (56 U12-U2 pairs and 4 U12-U12 pairs). In contrast, 29 of the 50 shared U12-type LECA introns in Pfams were paired with at least one other U12-type LECA intron in a paralog. The higher numbers of U12-U12 pairs in the Pfams set may result from multiple *bona fide* OGs being combined into a single KOG (see main text).

U12-type introns were less often in phase 0 and more in phase 2 than U2-type introns (supplementary fig. 6A, supplementary fig. 7A) and even more biased towards the 5' end (supplementary fig. 6B). Both observations are consistent with previous studies comparing intron types in present-day eukaryotes (Basu et al. 2008; Moyer et al. 2020). Assuming that nearly all type conversions were from U12 to U2 conversion, as has been argued based on comparative analyses (Sharp and Burge 1997), we inferred 32 U12 gains, 9 complete U12 losses and 7 U12-to-U2 conversions before duplications and a further 113 U12 gains, 20 complete U12 losses and 15 U12-to-U2 conversions on the branches that resulted in the LECA families. Paralogs that had a U12-type intron traced back to their pre-duplication lineage were overrepresented in cell cycle and inorganic ion transport and metabolism functions (supplementary fig. 7B). Differences in the fraction of U12-type introns among shared introns between functions of KOGs were not significant (supplementary fig. 6C) and only a few significant differences between different phylogenetic origins of these paralogs were found (supplementary fig. 7C). U12-type introns emerged at least before a large part of the complexification of the cell cycle and comparisons of the branch lengths seemed to suggest that U12-type introns are as old as U2-type introns, if not older (supplementary fig. 5B).

The three models have different expectations regarding shared U12-type introns, depending on the timing of minor intron emergence. The occurrence of both types among shared introns and the inferred age of U12-type introns are not consistent with the model of two "invasions" by group II introns that were clearly separated in time. Divergence from primordial introns in either separate lineages followed by fusion of these lineages or divergence in the same lineage is a more likely scenario based on our findings.

### Supplementary References

Basu MK, Makalowski W, Rogozin IB, Koonin EV. 2008. U12 intron positions are more strongly conserved between animals and plants than U2 intron positions. *Biol. Direct* 3:19.

Burge CB, Padgett RA, Sharp PA. 1998. Evolutionary fates and origins of U12-type introns. *Mol. Cell* 2:773–785.

Sharp PA, Burge CB. 1997. Classification of introns: U2-type or U12-type. *Cell* 91:875–879.

### Supplementary Figures

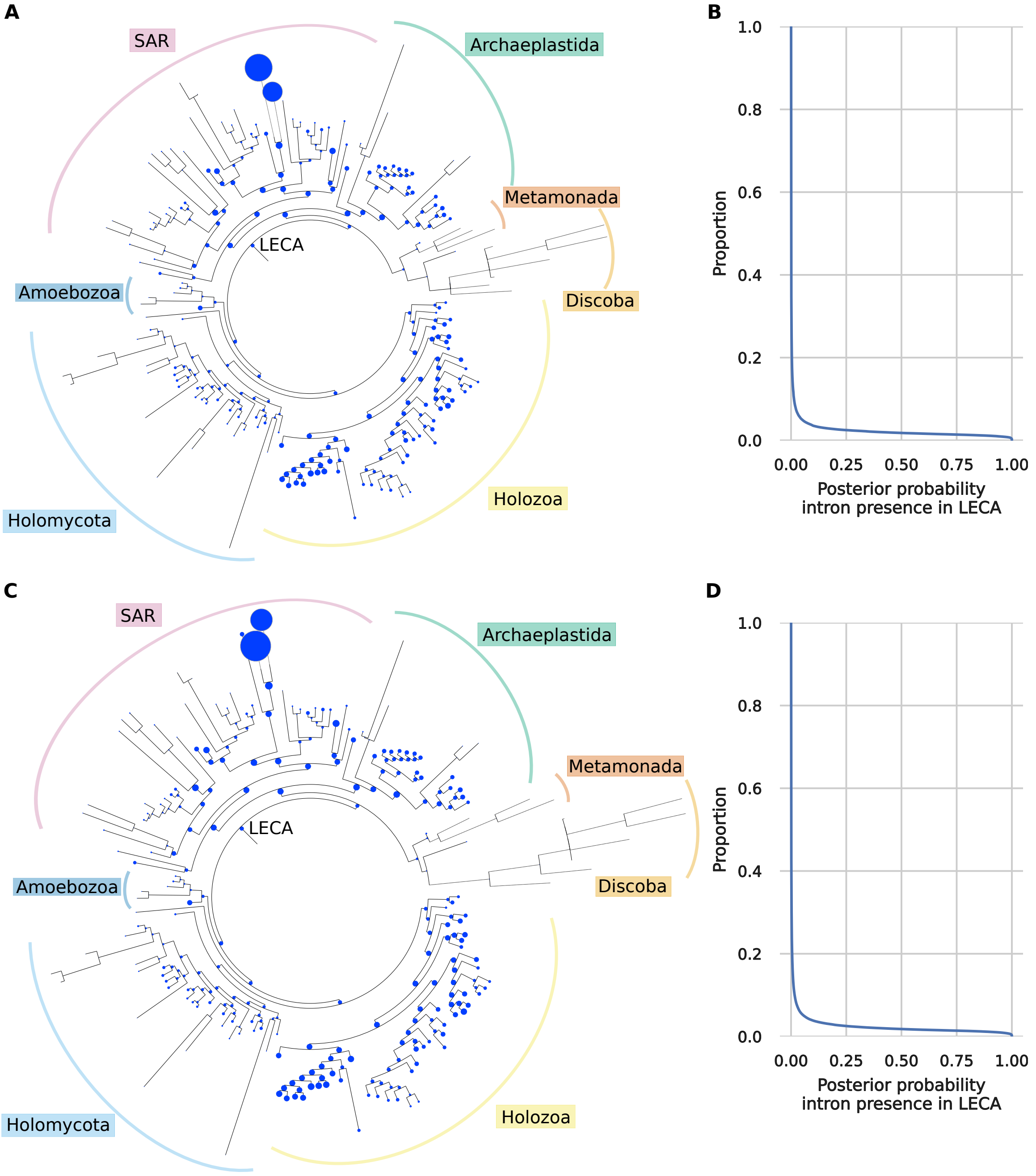


**Fig. S1. Ancestral intron reconstructions.** (*A*) Species tree with for each node the estimated number of introns (including missing sites) in KOGs represented as circles. These estimates are based on all used KOGs, including those from separate acquisitions. The size of the LECA node corresponds to 27108 introns. The two terminal nodes with the highest number of introns correspond to two dinoflagellate species. (*B*) Descending empirical cumulative distribution function plot of the posterior probability of a KOG intron to have been present in LECA. (*C*) Species tree with for each node the estimated number of introns (including missing sites) in Pfam OGs represented as circles. These estimates are based on all used Pfam OGs, including those from separate acquisitions. The size of the LECA node corresponds to 14977 introns. (*D*) Descending empirical cumulative distribution function plot of the posterior probability of a Pfam OG intron to have been present in LECA.

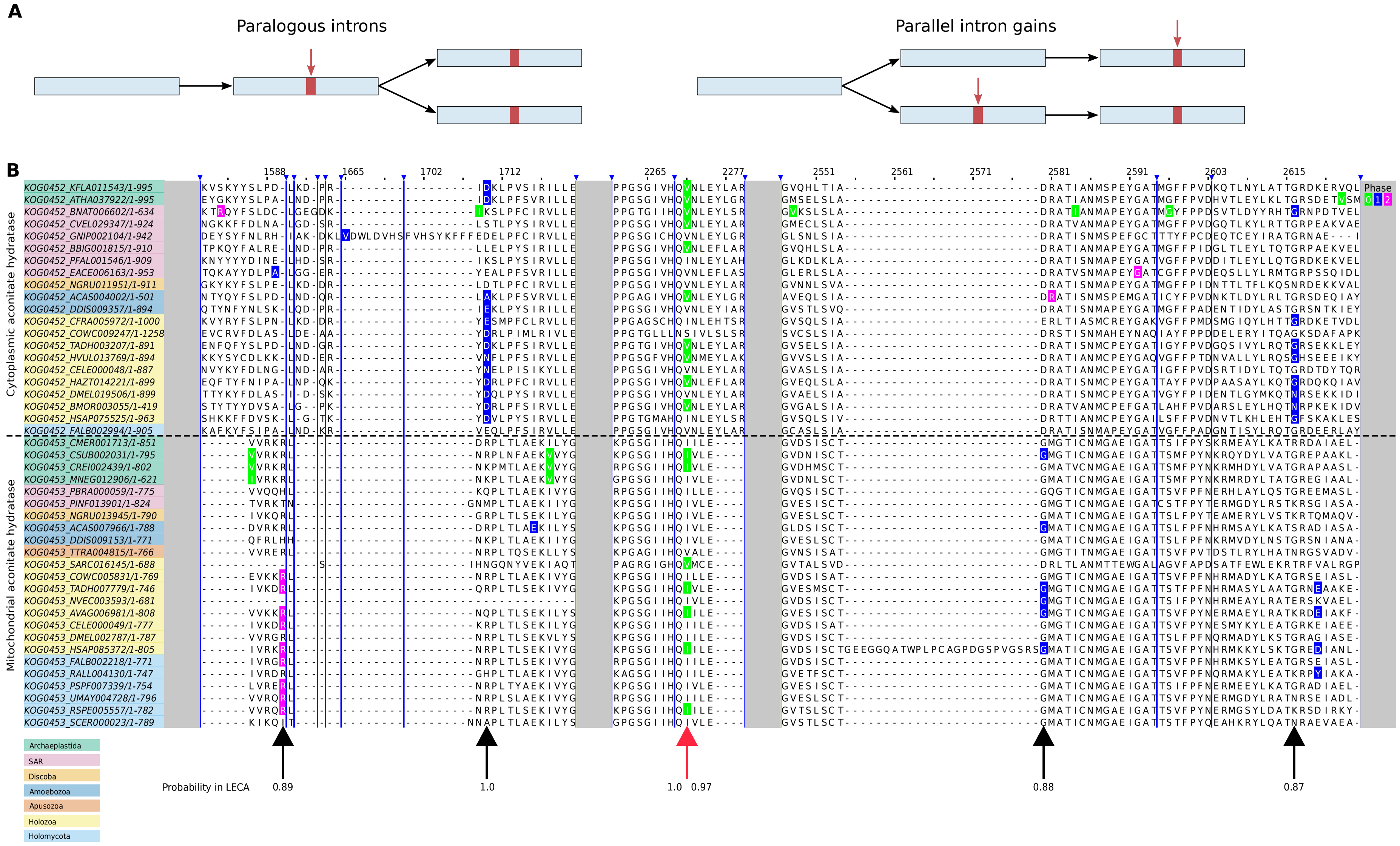
A
Paralogous introns
Parallel intron gains
B
Cytoplasmic aconitate hydratase
Mitochondrial aconitate hydratase
Phase
0 1 2
Archaeplastida
SAR
Discoba
Amoebozoa
Apusozoa
Holozoa
Holomycota
Probability in LECA
0.89
1.0
1.0 0.97
0.88
0.87

**Fig. S2. Parallel intron gains.** (*A*) Intron positions that are shared between paralogs could represent paralogous introns or could be due to parallel intron gains. (*B*) Example of parallel intron gains in separate acquisitions. Several aligned sequences from KOG0452 (cytoplasmic aconitate hydratase) and KOG0453 (mitochondrial aconitate hydratase) are shown with their mapped introns. The phase of introns is indicated with colours. Arrows point to introns that were probably present in LECA, with the number corresponding to the posterior probability. The shared intron that has been the result of parallel intron gains is indicated with a red arrow. Different sections of the alignment are separated by grey blocks and the blue stripes correspond to blocks of alignment positions that are only gaps in the sequences shown.

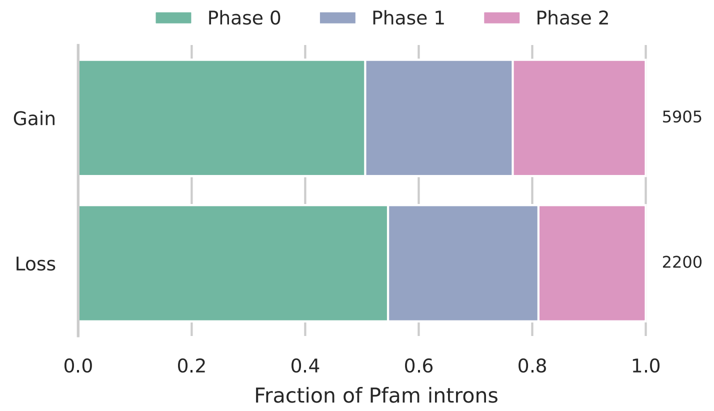


**Fig. S3. Phase distributions of introns in Pfams that were gained or lost before LECA.** Numbers indicate the number of inferred intron gains and losses.

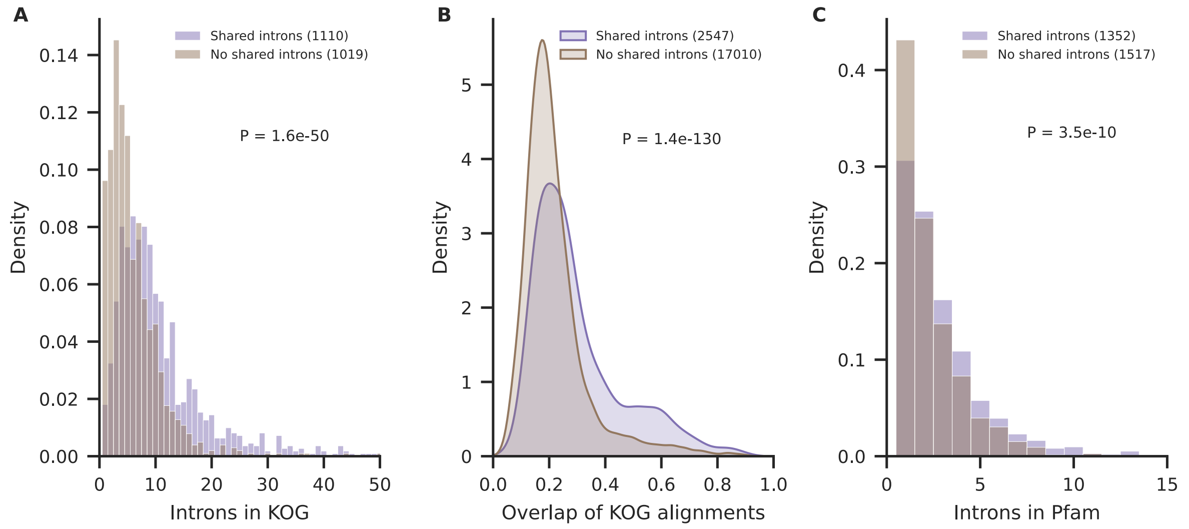


**Fig. S4. Influence of number of LECA introns and overlap of OG alignments on finding shared introns.** (*A*) Normalised histograms showing the distribution of the number of LECA introns in a KOG for KOGs with and without shared introns. For clarity, KOGs with more than 50 LECA introns are not shown. (*B*) Density plots showing the distribution of the fraction of overlapping positions in the alignment of all pairs of KOGs in the same cluster. Pairs with and without shared introns are depicted separately. Sites with more than 90% gaps were excluded in calculating the overlapping fraction. A lower overlap could be due to domain accretion and loss after duplication. (*C*) Normalised histograms showing the distribution of the number of LECA introns in a Pfam OG for Pfam OGs with and without shared introns. For clarity, Pfam OGs with more than 15 LECA introns are not shown. *P* values of Kolmogorov-Smirnov tests are shown. The numbers indicate the number of OGs (*A*, *C*) or pairs of KOGs (*B*). KOGs and Pfam OGs with no LECA introns were not included.

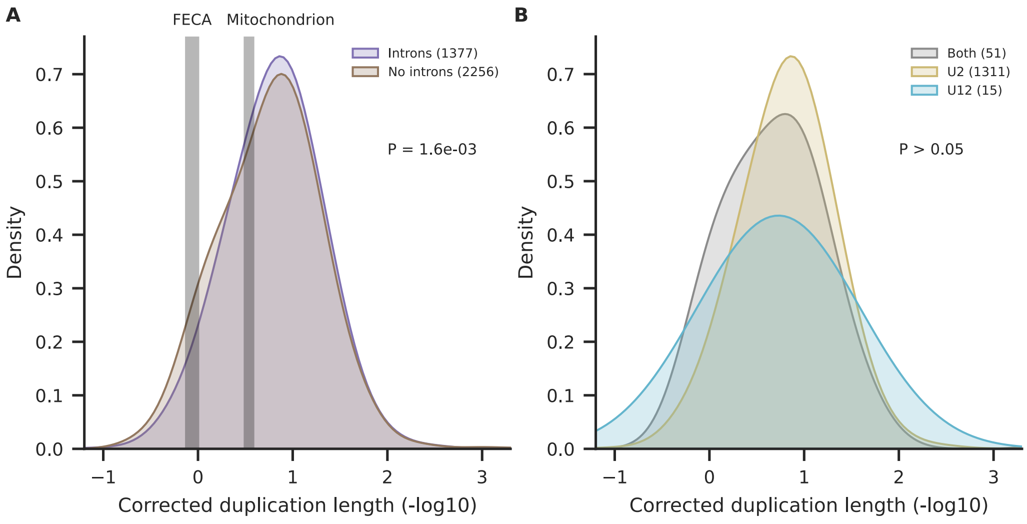


**Fig. S5. Timing of duplications with introns using branch lengths from phylogenetic trees.** (*A*) Density plot showing the duplication lengths of duplications with and without pre-duplication introns. For comparative purposes, the estimated timing (Vosseberg et al. 2021) of the first eukaryotic common ancestor (‘FECA’), which represents the divergence from the Asgard archaeal lineage, and the divergence from the alphaproteobacterial lineage (‘Mitochondrion’) are depicted in grey. The two distributions are significantly different according to the Kolmogorov-Smirnov test. (*B*) Density plot showing the duplication lengths of duplications with only U2-type, only U12-type or both types of pre-duplication introns. All pairwise comparisons with Kolmogorov-Smirnov tests were not significant. Numbers in both panels indicate the number of duplications considered.

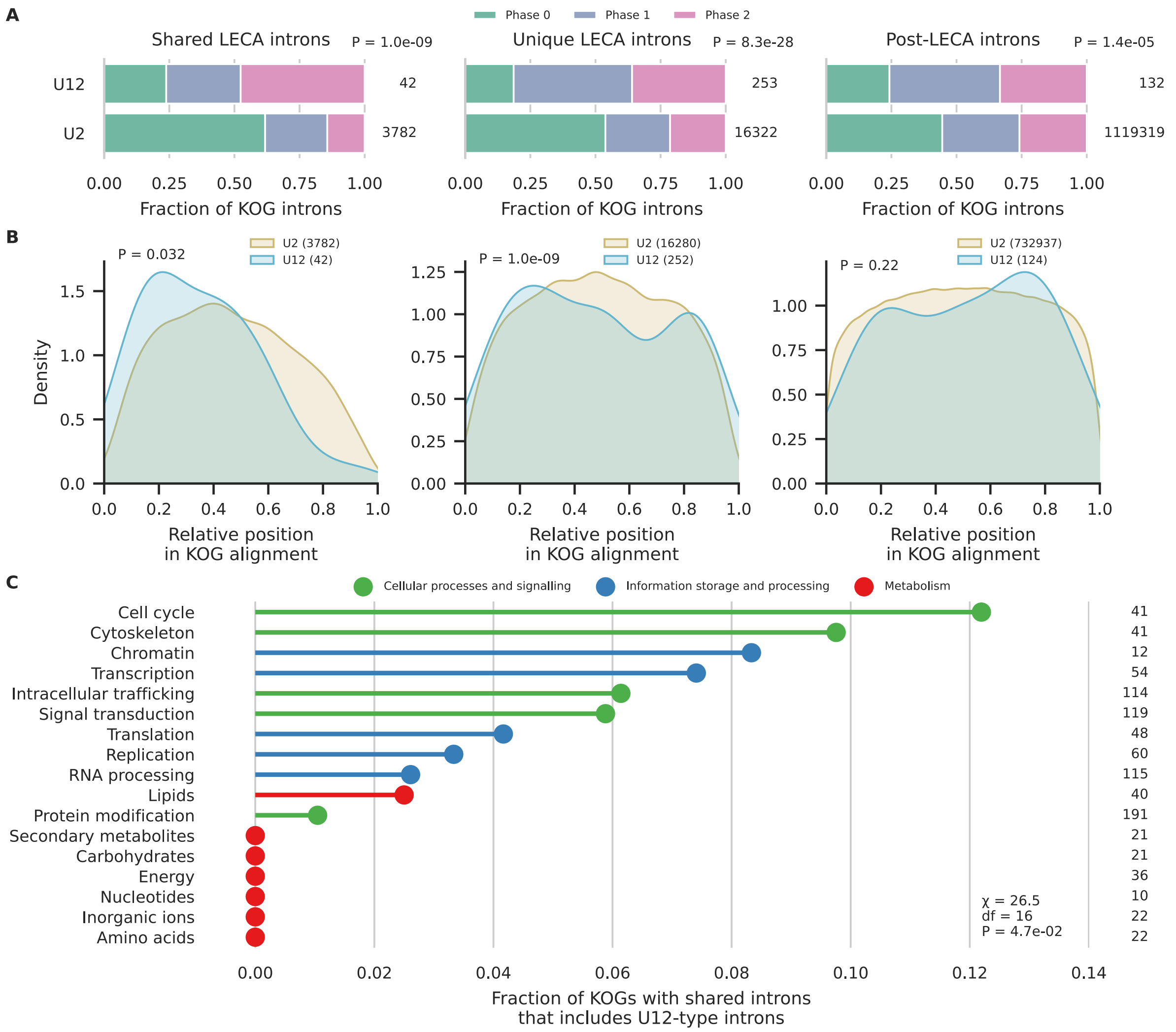


**Fig. S6. Comparison of U2- and U12-type introns in KOGs.** (*A*) Intron phase distributions for U2- and U12-type post-LECA, unique LECA and shared LECA introns in KOGs. *P* values of $\chi^2$ contingency tests are shown. (*B*) Density plots showing the relative positions of introns in the alignment of a KOG, comparing U2- and U12-type introns for the three different groups. *P* values of Kolmogorov-Smirnov tests are shown. (*C*) Fraction of KOGs with shared introns that includes U12-type introns in the different functional categories. Only functions with at least ten KOGs with shared introns are shown. Comparisons of the three functional categories (supplementary table 13) and pairwise comparisons of the different functions (supplementary table 14) were not significant. Numbers indicate the number of introns considered (*A*, *B*) or the number of KOGs with shared introns (*C*).

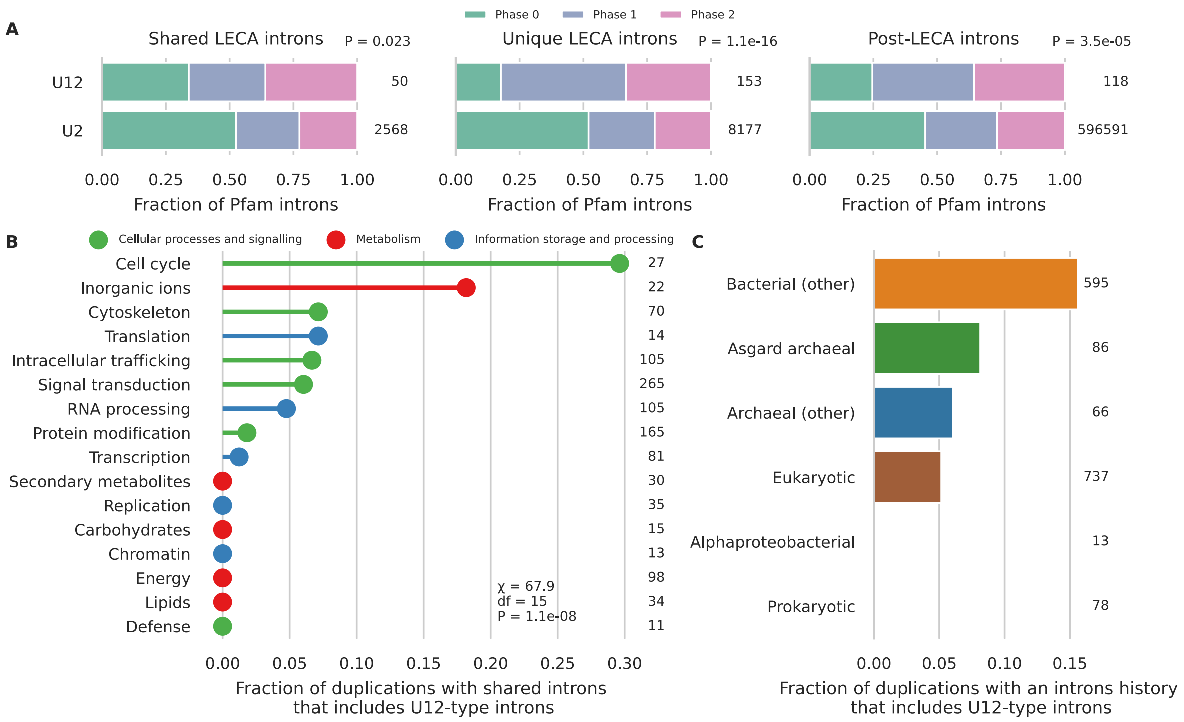


**Fig. S7. Comparison of U2- and U12-type introns in Pfams.** (*A*) Intron phase distributions for U2- and U12-type post-LECA, unique LECA and shared LECA introns in Pfam OGs. *P* values of $\chi^2$ contingency tests are shown. (*B*) Fraction of Pfam duplications with pre-duplication introns that includes U12-type introns in the different functional categories. Only functions with at least ten duplications with shared introns are shown. Comparisons of the three functional categories were not significant (supplementary table 15). 9.2% of pairwise comparisons were significant, which were only comparisons including the cell cycle and inorganic ions (supplementary table 16). (*C*) Fraction of Pfam duplications with pre-duplication introns in either that duplication or a more ancestral duplication that includes U12-type introns according to the different phylogenetic origins. Only the pairwise comparisons of bacterial and eukaryotic and bacterial and prokaryotic duplications were significant (supplementary table 17). Numbers indicate the number of introns (*A*) or the number of duplications (*B*, *C*) considered.

### Supplementary Tables

**Supplementary Table 1. KOG clusters.**

**Supplementary Table 2. Species list with links to used genome annotation files.**

**Supplementary Tables 3-17. Test statistics and *P* values of pairwise comparisons.**
